# Effects of *Arachis hypogaea* extract on TRPV4 activation and epidermal barrier function

**DOI:** 10.64898/2026.08.17.745342

**Authors:** Miyu Akiyama, Sayaka Takagi, Aya Yoshikoshi, Mari Iwase, Chihiro Honda, Tomoki Sato, Makoto Tominaga, Hisayoshi Hayashi, Shinji Miura, Shigenori Kumazawa, Kunitoshi Uchida

**Affiliations:** Laboratory of Functional Physiology, School of Food and Nutritional Sciences, University of Shizuoka, Yada 52-1, Suruga-ku, Shizuoka 422-8526 Japan; Graduate School of Integrated Pharmaceutical and Nutritional Sciences, University of Shizuoka, Yada 52-1, Suruga-ku, Shizuoka 422-8526 Japan; Laboratory of Food Analytical Chemistry, School of Food and Nutritional Sciences, University of Shizuoka, Yada 52-1, Suruga-ku, Shizuoka 422-8526 Japan; Laboratory of Nutritional Biochemistry, School of Food and Nutritional Sciences, University of Shizuoka, Yada 52-1, Suruga-ku, Shizuoka 422-8526 Japan; Thermal Biology Research Group, Nagoya Advanced Research and Development Center, Nagoya City University, Kawasumi 1, Mizuho-cho, Mizuho-ku, Nagoya 467-8601 Japan; Laboratory of Physiology, School of Food and Nutritional Sciences, University of Shizuoka, Yada 52-1, Suruga-ku, Shizuoka 422-8526 Japan

**Keywords:** TRPV4, *Arachis hypogaea* extract, epithelial barrier function, transepithelial electrical resistance

## Abstract

Transient receptor potential vanilloid 4 (TRPV4) is a Ca^2+^-permeable non-selective cation channel and its activating stimuli include anandamide, bisandrographolide, citric acid, arachidonic acid metabolic products by epoxygenases, hypo-osmotic cell swelling, and warm temperature. TRPV4 is involved in Ca^2+^-dependent signal transduction in several tissues. Since the activation of TRPV4 facilitates adherens junction formation in the skin epithelium, compounds that activate TRPV4 are expected to maintain or improve the barrier function of epidermal cells. In this study, we found that the extract of *Arachis hypogaea* (*A. hypogaea*) activate human TRPV4 (hTRPV4). In the Ca^2+^-imaging experiment, the application of *A. hypogaea* extract exhibited an increase in intracellular Ca^2+^ concentration ([Ca^2+^]_i_) in HEK293T cells expressing hTRPV4. The [Ca^2+^]_i_ increases by application of *A. hypogaea* extract were not observed in HEK293T cells expressing hTRPV1, mouse TRPV2, hTRPV3, hTRPM8, or hTRPA1. We then examined the physicochemical properties of the components responsible for TRPV4 activation. Ethanol extracts of A. hypogaea caused an increase in [Ca^2+^]_i_ in hTRPV4-expressing HEK293JN cells, whereas water, chloroform, and hexane extracts showed no activity. Moreover, the application of *A. hypogaea* extract enhanced transepithelial electrical resistance in the keratinocyte monolayer. These results suggest that *A. hypogaea* extract may contribute to the maintenance and improvement of the epidermal barrier function.

## INTRODUCTION

Most transient receptor potential (TRP) channels are Ca²⁺-permeable non-selective cation channels. In mammals, the TRP channel superfamily comprises 28 members classified into six subfamilies: TRPC (canonical), TRPM (melastatin), TRPML (mucolipin), TRPP (polycystin), TRPV (vanilloid), and TRPA (ankyrin). TRP channel activity is modulated by a variety of stimuli, including physical stimuli such as mechanical force and temperature, endogenous substances, ions, and lipids. TRP channels can also be activated or inhibited by exogenous chemical compounds. Physiologically, these channels contribute to sensory perception, intracellular signaling, and the maintenance of ion homeostasis (1, 2). TRP channels have also been implicated in various diseases, prompting extensive efforts to identify compounds that activate or inhibit these channels and to develop TRP channel-targeted therapeutics (3).

Similar to many other TRP channels, TRPV4 is a Ca²⁺-permeable non-selective cation channel. TRPV4 is widely expressed in tissues and cell types, including epithelial cells, the central nervous system, kidneys, urinary bladder, lungs, and adipose tissues. It is activated by membrane stretch associated with hypotonicity-induced cell swelling and by endogenous substances such as 5,6-epoxyeicosatrienoic acid (5,6-EET), a metabolite of arachidonic acid (4, 5). TRPV4 is also thermosensitive and has been reported to become activated at temperatures above approximately 27– 35°C (6). TRPV4 contributes to the maintenance of neuronal excitability in the brain (7), the regulation of urinary storage and voiding (8, 9), and the regulation of beige adipocyte differentiation (10). Thus, TRPV4 has diverse physiological functions in multiple tissues.

In addition to hypotonicity and temperature, TRPV4 is activated by several chemical compounds, including 4α-phorbol 12,13-didecanoate (4α-PDD), 2-aminoethoxydiphenyl borate (2-APB), anandamide, and citric acid. In contrast, TRPV4 activity is inhibited by ruthenium red, La³⁺, and Gd³⁺. Selective pharmacological modulators of TRPV4 have also been developed, including GSK1016790A, a selective TRPV4 agonist, and the antagonists HC067047 and GSK2193874 (11).

Several plant-derived compounds have also been reported to modulate TRPV4 activity. Bisandrographolide A derived from *Andrographis paniculata* (12), apigenin, a plant-derived flavone (13), and leaf extracts of *Lagerstroemia speciosa* (14) activate TRPV4, whereas berberine, which is found in plants including *Coptis japonica*, inhibits TRPV4 activity (15). Nevertheless, compared with other TRP channels such as TRPV1 and TRPA1, relatively few TRPV4 modulators from natural sources have been identified.

The skin is the largest organ covering the body surface and serves as a barrier that protects the body from external insults, including chemical substances and ultraviolet radiation, while preventing excessive water loss. The skin also functions as a sensory organ that detects touch, pain, and temperature and contributes to thermoregulation by controlling heat dissipation. It consists of three major layers: the epidermis, dermis, and subcutaneous tissue. The epidermis is further divided into the stratum corneum, stratum granulosum, stratum spinosum, and stratum basale. During differentiation, epidermal keratinocytes migrate from the basal layer through the spinous and granular layers toward the skin surface. These cells establish tight junctions (TJs) and adherens junctions (AJs), intercellular adhesion structures that contribute to epidermal barrier function (16). TJs are composed of proteins such as claudins and occludins, and are predominantly localized in the granular layer of the epidermis (17). AJs are formed by the accumulation of α-catenin, β-catenin, and E-cadherin with the actin cytoskeleton, with E-cadherin mediating adhesion between adjacent cells. TJs regulate the paracellular transport of fluids and solutes between epidermal cells, whereas AJs maintain tissue structure and regulate TJ formation (16). Together with the hydrophobic barrier provided by the stratum corneum, these intercellular junctions establish the robust barrier function of the skin. Accordingly, disruption of the epidermal barrier is associated with various diseases and disorders. For example, impairment of barrier function in atopic dermatitis increases immune sensitivity to environmental antigens such as house dust (18). Abnormal localization of TJ proteins and disruption of the epidermal barrier have also been reported in psoriatic lesions (19). Therefore, maintenance of an intact epidermal barrier is important not only for skin health but also for overall health. TRPV4 is reported to be involved in the maintenance of epidermal barrier function. Activation of TRPV4 expressed in epidermal keratinocytes increases the intracellular Ca²⁺ concentration and activates Rho proteins. These events promote actin filament reorganization and the formation and maturation of AJs and TJs, thereby strengthening intercellular adhesion (14, 20). Compounds that modulate TRPV4 activity may therefore have potential applications in maintaining epidermal barrier function and promoting skin health.

Based on this background, we searched for plant-derived compounds that modulate TRPV4 activity, and found that *Arachis hypogaea (A. hypogaea)* extract activated TRPV4. In the present study, we evaluated the TRPV4-activating effect of *A. hypogaea* extract and characterized its pharmacological properties.

## MATERIALS & METHODS

### HEK293JN cells stably expressing hTRPV4

Human embryonic kidney 293JN (HEK293JN) cells stably expressing human TRPV4 (hTRPV4) were maintained in high-glucose Dulbecco’s modified Eagle’s medium (DMEM; FUJIFILM Wako Pure Chemical Corporation, Osaka, Japan) supplemented with 0.2 mg/mL G418 sulfate (FUJIFILM Wako Pure Chemical Corporation), 50 units/mL penicillin (Thermo Fisher Scientific, Waltham, MA, USA), 50 μg/mL streptomycin (Thermo Fisher Scientific), 2 mM L-glutamine (Thermo Fisher Scientific), and 10% heat-inactivated fetal bovine serum (Biowest, Nuaillé, France). Cells were maintained in 10-cm culture dishes (Falcon, Corning Inc., NY, USA) at 37°C in a humidified atmosphere containing 5% CO₂ and passaged every 3–4 days. For Ca²⁺-imaging experiments, cells were seeded one day before the experiment onto 35-mm glass-bottom dishes (Matsunami Glass Ind., Ltd., Osaka, Japan) or 12-mm-diameter coverslips (Matsunami Glass Ind., Ltd.) placed in 35-mm culture dishes.

### Reagents and preparation of A. hypogaea extracts

Freeze-dried *A. hypogaea* seed samples were provided by the Industrial Research Institute of Shizuoka Prefecture. The samples were ground using a mortar and a mixer. One hundred mg of the ground sample was mixed with 500 μL of the indicated solvent, vortexed, and centrifuged. The resulting supernatant was used as the *A. hypogaea* extract. GSK1016790A, HC067047, capsaicin, carvacrol, allyl isothiocyanate (AITC), menthol, ethanol, chloroform, hexane, and dimethyl sulfoxide (DMSO) were purchased from FUJIFILM Wako Pure Chemical Corporation. 2-aminoethoxydiphenyl borate (2-APB) was purchased from Sigma-Aldrich (St. Louis, MO, USA). GSK1016790A, HC067047, 2-APB, carvacrol, and AITC were dissolved in DMSO and diluted to the indicated concentrations before use. Capsaicin and menthol were dissolved in ethanol and diluted before use.

### Culture and transfection of HEK293T cells

HEK293T cells were maintained in high-glucose DMEM supplemented with 50 units/mL penicillin, 50 μg/mL streptomycin, 2 mM L-glutamine, and 10% heat-inactivated fetal bovine serum. Cells were transfected one day before Ca²⁺ imaging experiments. One microgram of hTRPV1/pcDNA3.1, mTRPV2/pcDNA3.1, hTRPV3/pcDNA3.1, hTRPV4/pcDNA3.1, hTRPA1/pcDNA3.1, or hTRPM8/pcDNA3.1 plasmid, together with 0.1 μg of a DsRed expression plasmid for identification of transfected cells, was added to 100 μL Opti-MEM (Thermo Fisher Scientific). Six microliters of PLUS Reagent (Thermo Fisher Scientific) was then added, and the mixture was incubated for 10–15 min. Separately, 4 μL Lipofectamine Reagent (Thermo Fisher Scientific) was added to 100 μL Opti-MEM. The two solutions were combined and incubated for an additional 10–15 min to prepare the transfection mixture. The culture medium was removed from the HEK293T cells, and 800 μL Opti-MEM and the transfection mixture were added. Cells were incubated at 37°C under 5% CO₂. After 3–4 h, the cells were reseeded onto 35-mm glass-bottom dishes and used for experiments one day after transfection.

### Ca²⁺-imaging

Cells were loaded with 3.3 μM Fluo-4 AM (Dojindo Laboratories, Kumamoto, Japan) for at least 40 min. For experiments using glass-bottom dishes, the culture medium was completely removed immediately before measurement. Cells were washed two or three times with bath solution containing 140 mM NaCl, 5 mM KCl, 2 mM MgCl₂, 2 mM CaCl₂, 10 mM HEPES, and 10 mM glucose, adjusted to pH 7.4. One milliliter of fresh bath solution was then added to the dish. At 30 s after the start of recording, 500 μL bath solution containing the sample extract was added. At 90 s, 500 μL bath solution containing an agonist for the respective TRP channel was added. At 180 s, Ionomycin was added at a final concentration of 3 μM. For inhibition experiments, HC067047, a TRPV4 antagonist was added immediately before recording at a final concentration of 10 μM. For experiments using a perfusion system, coverslips were placed in an open recording chamber (Warner Instruments, Hamden, CT, USA), and bath solution was perfused at a flow rate of 5 mL/min. Solutions were switched using an electronically controlled valve system (VC-6; Warner Instruments). Fluorescence images were acquired every 5 s using an inverted fluorescence microscope (Ti2; Nikon Corporation, Tokyo, Japan) equipped with a CoolSNAP ES CCD camera (Photometrics, Tucson, AZ, USA). Fluorescence signals were recorded using NIS-Elements software (Nikon Corporation) and analyzed using ImageJ software (National Institutes of Health, Bethesda, MD, USA).

### Culture of HaCaT cells

HaCaT human epidermal keratinocytes were maintained at 37°C in a humidified atmosphere containing 5% CO₂. Cells were cultured in calcium-free DMEM (Nacalai Tesque, Inc., Kyoto, Japan) supplemented with 10% heat-inactivated, calcium-depleted fetal bovine serum, 0.04 mM CaCl₂, 100 units/mL penicillin, 100 μg/mL streptomycin, and 2 mM L-glutamine. The culture medium was replaced every 2–3 days. Cells were passaged every 4–6 days by detachment from the culture dishes using Accutase (Funakoshi Co., Ltd., Tokyo, Japan).

### Measurement of transepithelial electrical resistance

HaCaT cells were seeded at 1 × 10⁵ cells/mL onto 12-well cell culture inserts with a pore size of 0.4 μm (Falcon, Corning Inc., NY, USA). Five hundred μL of culture medium was added to the insert, and 1.5 mL was added to the lower chamber. Cells were cultured for 2 days. Transepithelial electrical resistance (TEER) was measured using a Millicell ERS-2 Electrical Resistance System (EMD Millipore Corporation, Billerica, MA, USA). After the initial measurement, the culture medium was replaced with DMEM containing 1 mM CaCl₂, with or without 0.1% (v/v) *A. hypogaea* extract. TEER was measured again 1 and 2 days after replacement of the medium.

### Statistical analysis

Data are presented as the mean ± standard error of the mean (SEM). Comparisons between two groups were performed using Student’s *t*-test. A value of p < 0.05 was considered statistically significant.

## Results

### Activation of TRPV4 by A. hypogaea extract

To examine the effect of *A. hypogaea* extract on human TRPV4 (hTRPV4), *A. hypogaea* extract diluted 1:1,000 in bath solution was applied to HEK293JN cells stably expressing hTRPV4. Application of the *A. hypogaea* extract induced an increase in intracellular Ca²⁺ concentration ([Ca^2+^]_i_) (Fig. 1A). We next applied the *A. hypogaea* extract to hTRPV4-expressing HEK293JN cells pretreated with 10 μM HC067047, a TRPV4 antagonist. Under these conditions, the *A. hypogaea* extract-induced increase in [Ca^2+^]_i_ was almost completely abolished (Fig. 1B). The response to a high concentration of the TRPV4 agonist (30 μM GSK1016790A) was also attenuated in the presence of HC067047, confirming the inhibitory effect of HC067047 on TRPV4 activity. The *A. hypogaea* extract-induced Ca²⁺ response was quantified as the ratio of the peak increase induced by the *A. hypogaea* extract to the peak increase induced by Ionomycin. The relative increase in [Ca^2+^]_i_ induced by the *A. hypogaea* extract was 5.32 ± 0.90% in the presence of HC067047, which was significantly lower than that observed in the absence of HC067047 (35.81 ± 3.11%; Fig. 1C).

**Figure 1.**
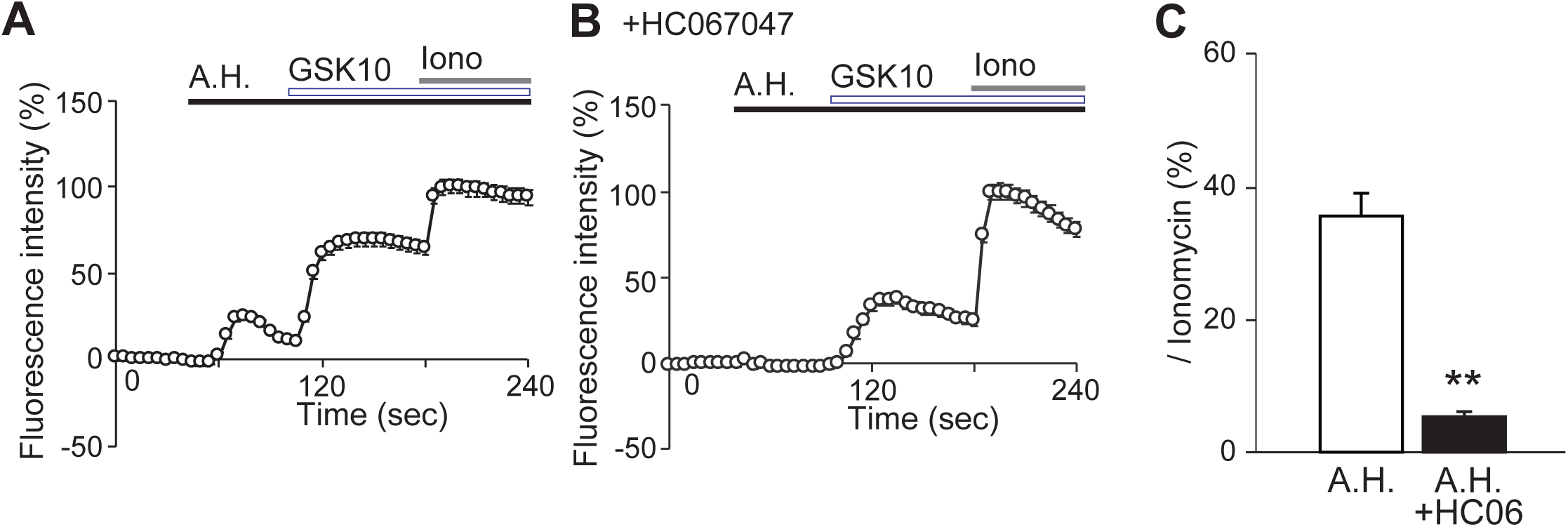
Activation of TRPV4 by *Arachis hypogaea* extract. (A) Time course of intracellular Ca^2+^ concentration ([Ca^2+^]_i_) changes in HEK293JN cells stably expressing hTRPV4 during sequential application of *A. hypogaea* extract diluted 1:1,000 (A.H.), GSK1016790A (3 μM; GSK10), and Ionomycin (2 μM; Iono). (B) Time course of intracellular Ca^2+^ concentration ([Ca^2+^]_i_) changes in stably hTRPV4-expressing HEK293JN cells pretreated with HC067047, a TRPV4 antagonist (10 μM) immediately before recording, followed by sequential application of *A. hypogaea* extract diluted 1:1,000 (A.H.), GSK1016790A (30 μM; GSK10), and Ionomycin (2 μM; Iono). (C) Quantification of [Ca^2+^]_i_ increases by treatment of *A. hypogaea* extract, calculated as the peak increase in [Ca^2+^]_i_ induced by the extract divided by the peak increase induced by Ionomycin. Data are presented as the mean ± SEM; n = 121-130. \*\**p* < 0.01, Student’s *t*-test.

We next examined the concentration dependence of the effect of *A. hypogaea* extract on hTRPV4 activity. TRPV4 activity in response to *A. hypogaea* extract diluted 1:10,000, 1:3,000, 1:1,000, or 1:300 was evaluated using Ca²⁺-imaging. *A. hypogaea* extract diluted 1:10,000 or 1:3,000 produced little or no response (Fig. 2A, B), whereas the 1:1,000 and 1:300 dilutions induced concentration-dependent increases in [Ca^2+^]_i_ (Fig. 2C, D). In the continued presence of *A. hypogaea* extract diluted 1:300, application of the TRPV4 agonist decreased the [Ca^2+^]_i_, suggesting that a high concentration of *A. hypogaea* extract may attenuate the response to the TRPV4 agonist. Quantification of the Ca²⁺ responses showed that the relative [Ca^2+^]_i_ increases were 9.40 ± 1.57%, 5.42 ± 0.94%, 32.05 ± 2.35%, and 67.96 ± 7.26% at dilutions of 1:10,000, 1:3,000, 1:1,000, and 1:300, respectively (Fig. 2E).

**Figure 2.**
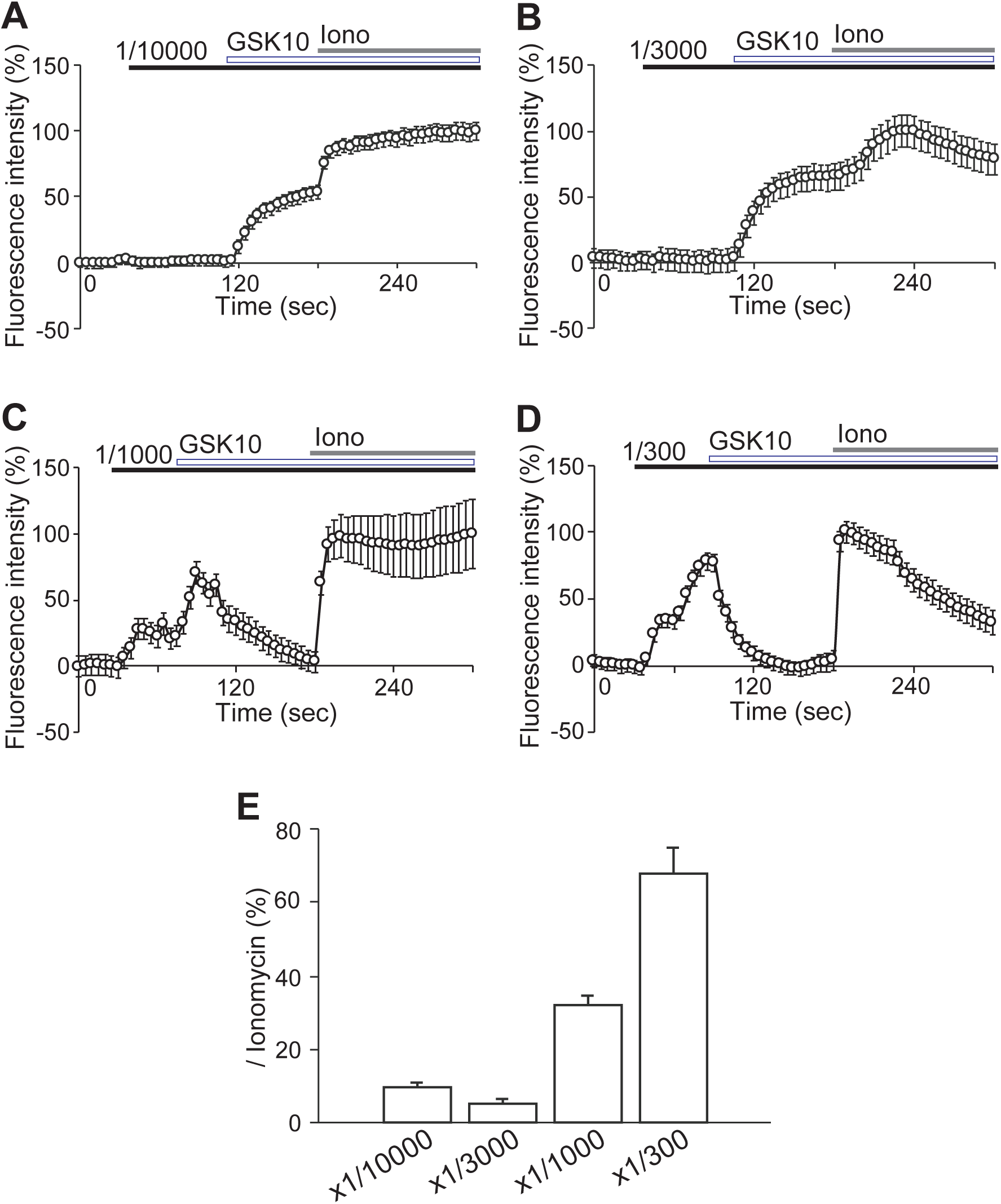
Concentration-dependent activation of TRPV4 by *Arachis hypogaea* extract. (A-D) Time course of intracellular Ca^2+^ concentration ([Ca^2+^]_i_) changes in HEK293JN cells stably expressing hTRPV4 during sequential application of *A. hypogaea* extract diluted 1:10,000 (A), 1:3,000 (B), 1:1,000 (C), or 1:300 (D), followed by GSK1016790A (3 μM; GSK10), and Ionomycin (2 μM; Iono). (E) Quantification of [Ca^2+^]_i_ increases by treatment of *A. hypogaea* extract, calculated as the peak increase in [Ca^2+^]_i_ induced by the extract divided by the peak increase induced by Ionomycin. Data are presented as the mean ± SEM; n = 138-261.

### Effects of A. hypogaea extract on TRP channels predominantly expressed in sensory neurons and the epidermis

To examine the channel specificity of *A. hypogaea* extract, we tested its effects on hTRPV1, mouse TRPV2 (mTRPV2), hTRPV3, hTRPA1, and hTRPM8, which are expressed in sensory neurons and/or the epidermis. The effects of *A. hypogaea* extract were evaluated by Ca²⁺-imaging in HEK293T cells transiently expressing each TRP channel. *A. hypogaea* extract diluted 1:1,000 did not induce a detectable increase in [Ca^2+^]_i_ in cells expressing hTRPV1 (Fig. 3A), mTRPV2 (Fig. 3B), hTRPV3 (Fig. 3C), hTRPA1 (Fig. 3E), or hTRPM8 (Fig. 3F), whereas a clear response was observed in cells expressing hTRPV4 (Fig. 3D). Application of a representative agonist for each channel induced a marked increase in [Ca^2+^]_i_, confirming the functional expression of the respective TRP channels. *A. hypogaea* extract also failed to induce an increase in [Ca^2+^]_i_ in mock-transfected HEK293T cells (Fig. 3G). Quantification of the Ca²⁺ responses showed that the relative increase in [Ca^2+^]_i_ was 51.80 ± 3.46% in hTRPV4-expressing cells, whereas the responses in cells expressing each of the other five TRP channels were less than 10%.

**Figure 3.**
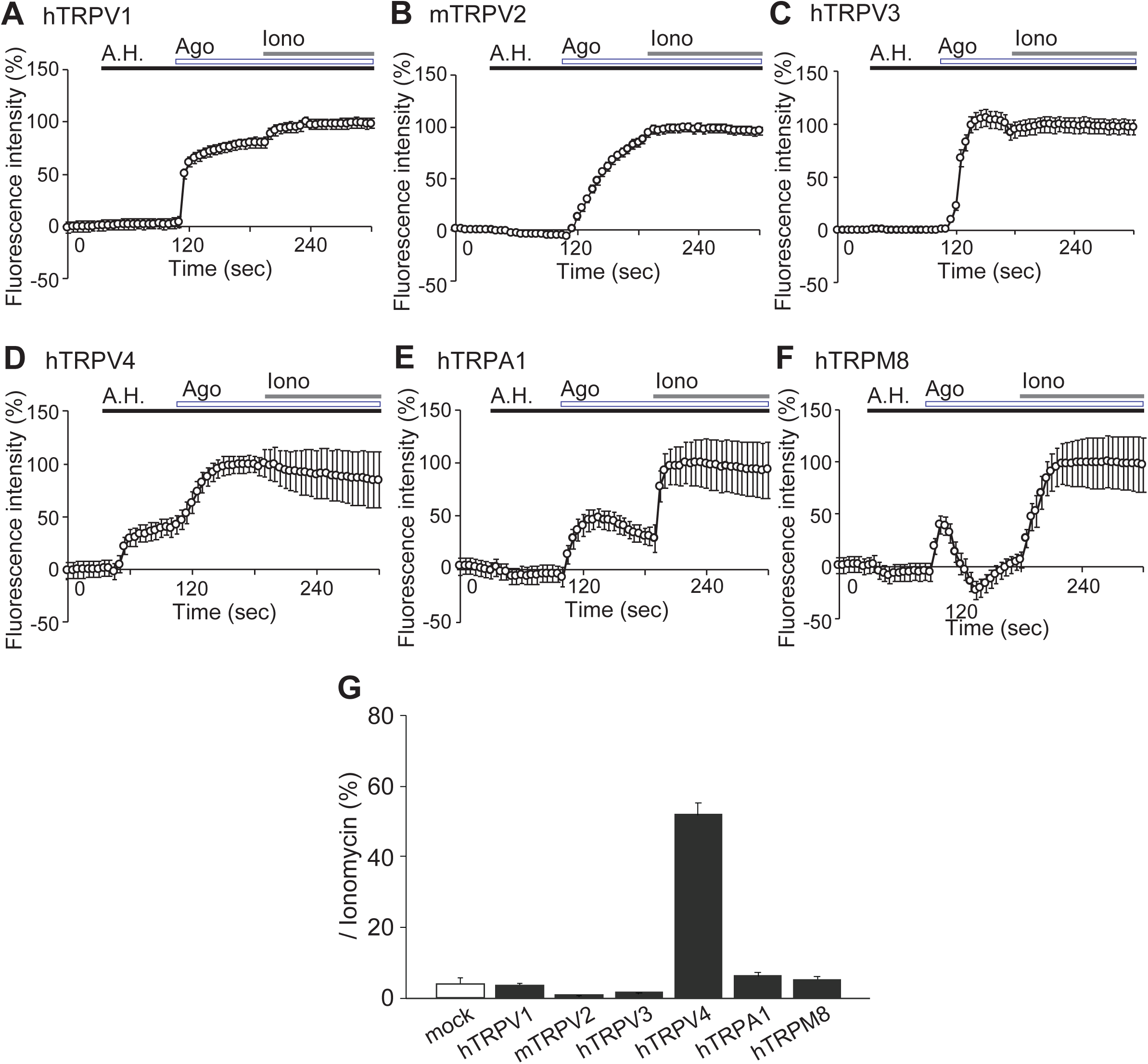
Effects of Arachis hypogaea extract on TRP channels expressed in sensory neurons and the skin. (A-F) Time course of intracellular Ca^2+^ concentration ([Ca^2+^]_i_) changes in HEK293T cells transiently expressing hTRPV1 (A), mTRPV2 (B), hTRPV3 (C), hTRPA1 (E), hTRPM8 (F), and in HEK293JN cells stably expressing hTRPV4 (D). *A. hypogaea* extract diluted 1:1,000 was applied, followed by a representative agonist for each TRP channel: capsaicin for hTRPV1 (1 μM), 2-APB for mTRPV2 (500 μM), 2-APB plus carvacrol for hTRPV3 (250 and 500 μM, respectively), GSK1016790A for hTRPV4 (3 μM), AITC for hTRPA1 (100 μM), or menthol for hTRPM8 (500 μM). Ionomycin (2 μM; Iono) was subsequently applied. (G) Quantification of [Ca^2+^]_i_ increases by treatment of *A. hypogaea* extract in mock-transfected cells and cells expressing the indicated TRP channels. Responses were calculated as the peak increase in [Ca^2+^]_i_ induced by the extract divided by the peak increase induced by Ionomycin. Data are presented as mean ± SEM; n = 48-406.

### Properties of the TRPV4-activating component in A. hypogaea extract

To characterize the TRPV4-activating component in *A. hypogaea* extract, we first examined whether its activity was affected by storage conditions. *A. hypogaea* DMSO extract was diluted in bath solution and stored at 4°C for 1 day before application to hTRPV4-expressing HEK293JN cells. The stored extract induced little or no increase in [Ca^2+^]_i_ (Fig. 4A, B). The relative increase in [Ca^2+^]_i_ was 66.54 ± 5.10% for freshly diluted *A. hypogaea* extract but only 4.42 ± 0.85% for the extract stored for 1 day after dilution (Fig. 4C).

**Figure 4.**
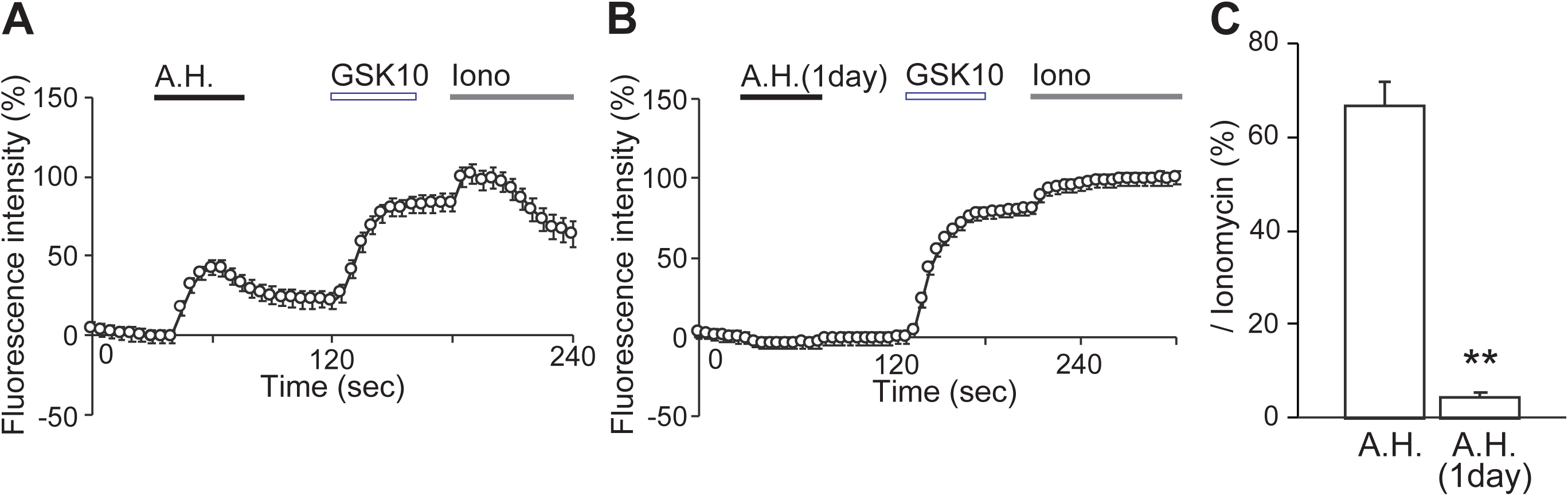
Stability of the TRPV4-activating component in *Arachis hypogaea* extract in aqueous solution. (A) Time course of intracellular Ca^2+^ concentration ([Ca^2+^]_i_) changes in HEK293JN cells stably expressing hTRPV4 during application of *A. hypogaea* extract diluted 1:1,000 in bath solution immediately before use (A) or stored for 1 day after dilution in bath solution (B), followed by GSK1016790A (3 μM; GSK10), and Ionomycin (2 μM; Iono). (C) Quantification of [Ca^2+^]_i_ increases by treatment of *A. hypogaea* extract, calculated as the peak increase in [Ca^2+^]_i_ induced by the extract divided by the peak increase induced by Ionomycin. Data are presented as the mean ± SEM; n = 83-84. \*\**p* < 0.01, Student’s *t*-test.

We next evaluated the effects of the extraction solvent on TRPV4-activating activity. *A. hypogaea* extracts were prepared using purified water, ethanol, chloroform, or hexane. The water, chloroform, and hexane extracts did not induce a detectable increase in [Ca^2+^]_i_, even at a dilution of 1:300. In contrast, the ethanol extract induced concentration-dependent [Ca^2+^]_i_ increases at dilutions of 1:1,000 and 1:300, although the responses were smaller than those induced by the DMSO extract (Fig. 5A–D). Among the extracts tested at a 1:300 dilution, the ethanol extract produced the largest relative increase in [Ca^2+^]_i_ (26.27 ± 1.86%), whereas only small or no responses were observed with the water (1.51 ± 1.50%), chloroform (3.81 ± 1.42%), and hexane extract (−0.62 ± 1.26%) (Fig. 5E).

**Figure 5.**
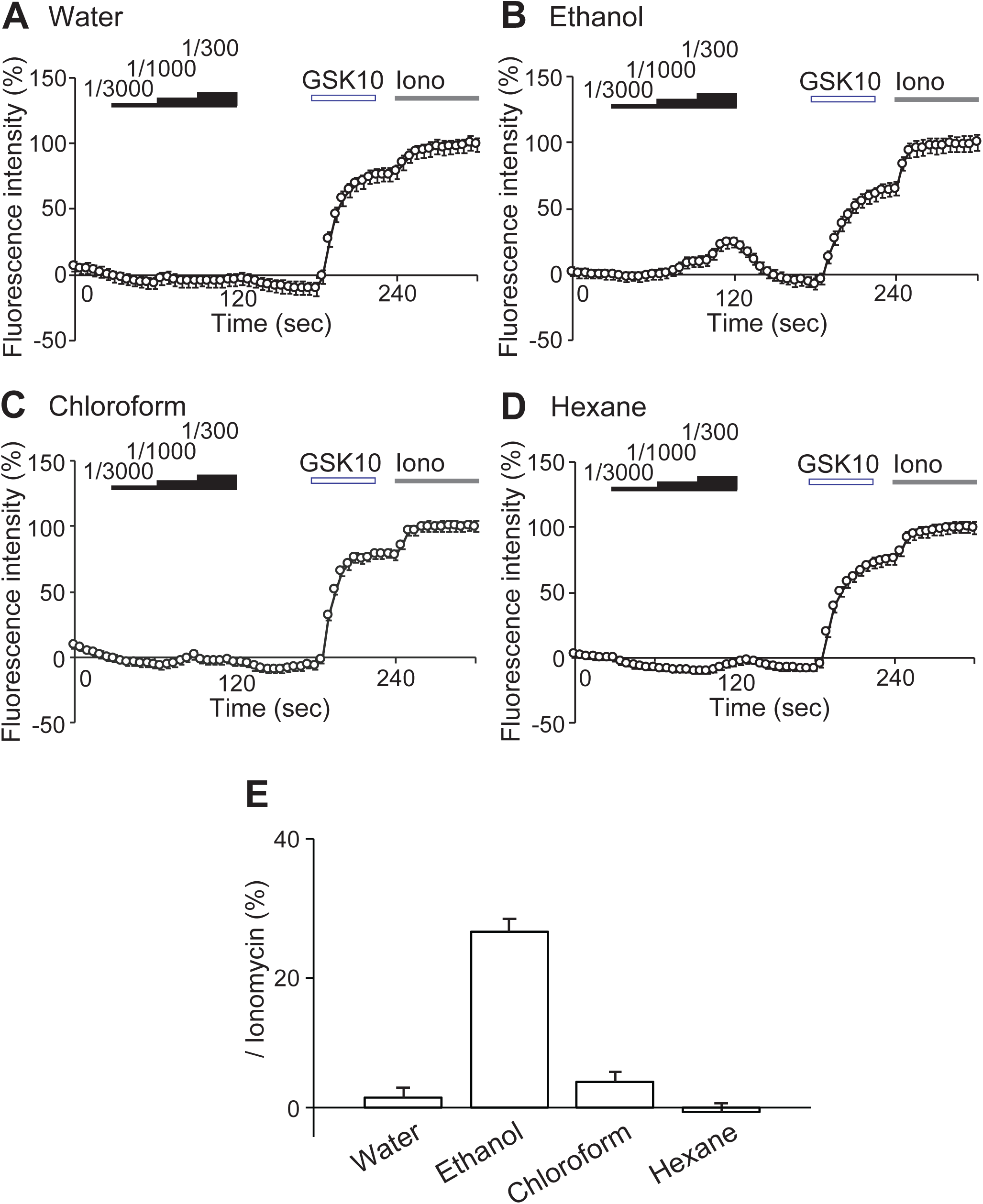
Physicochemical properties of the TRPV4-activating component in *Arachis hypogaea* extract. (A-D) Time course of intracellular Ca^2+^ concentration ([Ca^2+^]_i_) changes in HEK293JN cells stably expressing hTRPV4 during application of *A. hypogaea* extract prepared with purified water (A), ethanol (B), chloroform (C), or hexane (D). Each extract was tested at dilutions of 1:3,000, 1:1,000, and 1:300, followed by GSK1016790A (3 μM; GSK10), and Ionomycin (2 μM; Iono). (E) Quantification of [Ca^2+^]_i_ increases by treatment of *A. hypogaea* extract, calculated as the peak increase in [Ca^2+^]_i_ induced by the extract divided by the peak increase induced by Ionomycin. Data are presented as mean + SEM; n = 132-171.

### Effect of A. hypogaea extract on epidermal barrier function

To examine the effect of *A. hypogaea* on epidermal barrier function, trans-epithelial electrical resistance (TEER) was measured 1 and 2 days after the addition of 1 mM CaCl_2_ to confluent monolayers of HaCaT cells. TEER values were expressed relative to the value measured immediately before the addition of 1 mM CaCl₂, which was defined as 100%. The addition of 1 mM CaCl_2_ increased TEER in a time-dependent manner, reaching 110.9 ± 3.0% after 48 h. Cotreatment with *A. hypogaea* extract further increased TEER to 128.9 ± 5.1% after 48 h, compared with treatment with 1 mM CaCl_2_ alone.

## Discussion

Activation of TRPV4 in epidermal keratinocytes increases the intracellular Ca²⁺ concentration and activates downstream signaling molecules, including Rho-family GTPases. Activation of Rho signaling induces reorganization of the cortical actin cytoskeleton and promotes the accumulation of α-catenin, β-catenin, and E-cadherin along actin filaments, thereby facilitating the formation of E-cadherin-mediated intercellular adherens junctions (20). Through the promotion of intercellular junction formation, TRPV4 activation strengthens epidermal barrier function. Thus, compounds that activate TRPV4 may contribute to the maintenance or restoration of the skin barrier. In the present study, *A. hypogaea* extract not only activated TRPV4 but also increased TEER in epidermal keratinocytes (Figs. 1 and 6). In epidermal cells, a calcium switch induces the formation and maturation of intercellular junctions, and TEER increases as tight junction-dependent barrier function develops (21). Therefore, the increase in TEER induced by *A. hypogaea* extract could reflect enhanced epithelial barrier function resulting from the formation or maturation of intercellular junctions. However, the present study provides no direct evidence linking TRPV4 activation by A. hypogaea extract to the extract-induced TEER increase. Further studies are required to determine whether this effect is dependent on TRPV4 activation.

**Figure 6.**
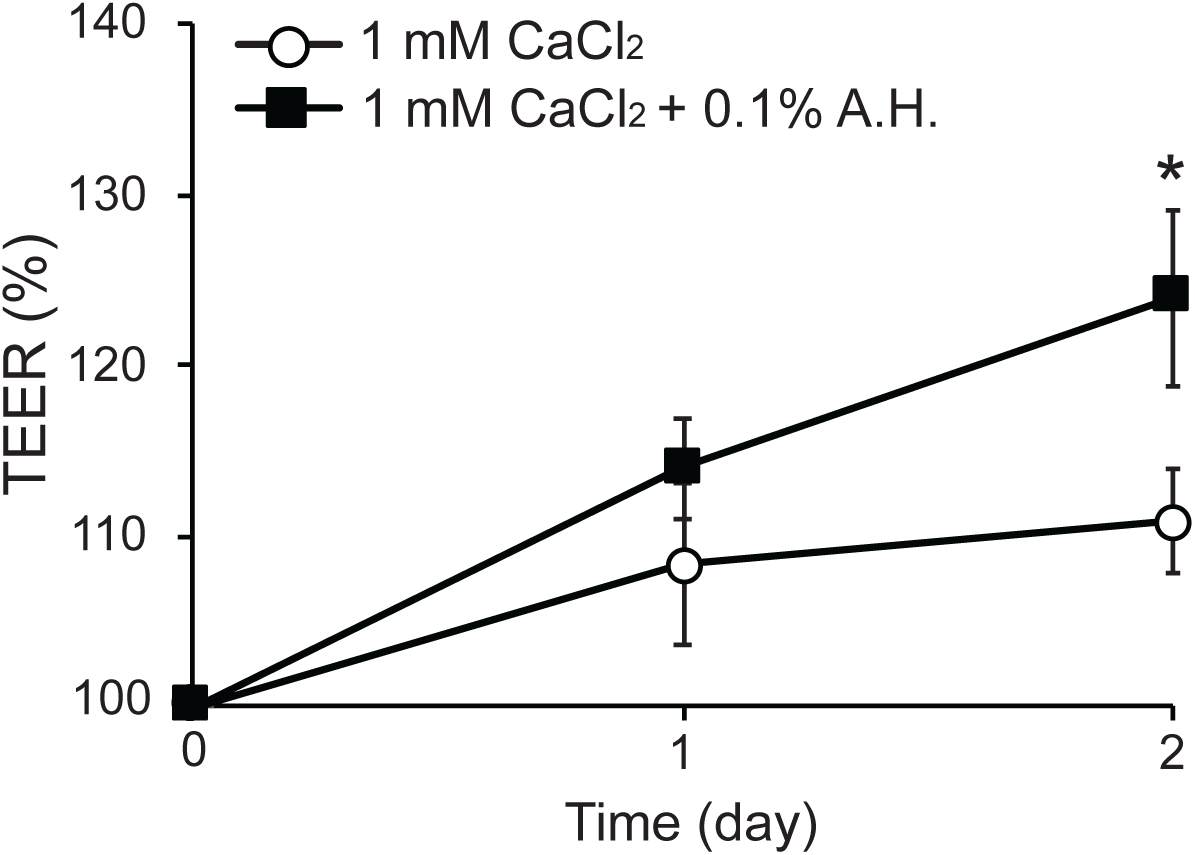
Effect of *Arachis hypogaea* extract on barrier function in HaCaT cells. Changes in trans-epithelial electrical resistance (TEER) in HaCaT cells after treatment with 1 mM CaCl₂ alone or 1 mM CaCl₂ together with 0.1% *A. hypogaea* extract. TEER values were normalized to the value measured immediately before each treatment, which was defined as 100%. Data are presented as the mean ± SEM; n = 6-7. *p < 0.05, Student’s *t*-test.

TRPV4 is expressed not only in epidermal keratinocytes but also in diverse tissues, including the central nervous system, kidneys, and adipose tissues (11). Its physiological effects are likely to vary depending on the tissue and the nature of the activating stimulus. Therefore, the effects of TRPV4-activating components in *A. hypogaea* extract on tissues other than the epidermis should be evaluated carefully. Nevertheless, *A. hypogaea* extract could exert physiological effects beyond the regulation of epidermal barrier function.

A. hypogaea extract did not produce a clear response in other TRP channels expressed in epidermal cells or sensory neurons. TRPV1 and TRPA1 are involved in pain and irritant sensations, TRPM8 mediates cool sensation (22), and TRPV3 has been associated with inflammatory skin diseases (23). The absence of detectable activation of these TRP channels by *A. hypogaea* extract may suggest that it is less likely to induce undesirable sensory effects when applied to the skin. However, actual skin irritation and safety must be evaluated directly. In particular, the possibility of allergic reactions to *A. hypogaea*-derived components requires careful investigation.

The TRPV4-activating activity observed in this study was present in fractions extracted with ethanol or DMSO but was not detected in extracts prepared with water, chloroform, or hexane. These solvent-dependent properties suggest that the active component is unlikely to be a highly hydrophobic lipid such as a triacylglycerol and may instead be a compound of intermediate polarity that is soluble in ethanol and DMSO. Amphiphilic lipids, including fatty acid derivatives and monoacylglycerols, can also dissolve in ethanol and DMSO and therefore remain possible candidates for the active component. The marked loss of activity after dilution and storage in aqueous solution further suggests that the active component is unlikely to be a stable, highly water-soluble small molecule. Its apparent effective concentration may decrease in aqueous solution because of degradation, oxidation, photodegradation, aggregation, or other physicochemical processes. In addition to lipids, *A. hypogaeas* contain several classes of plant secondary metabolites, including phenolic acids, flavonoids, and stilbenes (24). Stilbenes such as resveratrol are representative *A. hypogaea* polyphenols. However, because resveratrol has been reported to potentially inhibit TRPV4 activity (25), it is unlikely to be the major component responsible for the TRPV4 activation observed in the present study. *A. hypogaeas* also contain catechins, procyanidins, and *p*-coumaric acid derivatives (24). These plant secondary metabolites, amphiphilic lipids, or unidentified compounds of intermediate polarity could contribute to the activation of TRPV4. Further fractionation of the ethanol-soluble fraction and identification of its active constituent or constituents will be necessary to clarify the mechanism by which *A. hypogaea* extract activates TRPV4.

## Acknowledgments

We thank the members of the Food Science Section, Industrial Research Institute of Shizuoka Prefecture, for providing the freeze-dried *A. hypogaeas* samples.

## Author contributions

K.U. designed the experiments. M.A., S.T., and K.U. wrote the manuscript. M.A., S.T., A.Y. C.H., T.S., S.K., and K.U. performed the experiments and analyzed the data. M.I., C.H., T.S., M.T., H.H., S.M., S.K., and K.U. discussed and interpreted the data. All authors read and approved the final manuscript.

## Abbreviations

2-APB: 2-aminoethoxydiphenyl borate
5,6-EET: 5,6-epoxyeicosatrienoic acid
AITC: allyl isothiocyanate
AJ: adherens junction
Ca²?: calcium ion
DMEM: Dulbecco’s modified Eagle’s medium
DMSO: dimethyl sulfoxide
FBS: fetal bovine serum
HEK: human embryonic kidney
HEPES: 4-(2-hydroxyethyl)-1-piperazineethanesulfonic acid
TEER: transepithelial electrical resistance
TJ: tight junction
TRP: transient receptor potential
TRPA: transient receptor potential ankyrin
TRPC: transient receptor potential canonical
TRPM: transient receptor potential melastatin
TRPML: transient receptor potential mucolipin
TRPP: transient receptor potential polycystin
TRPV: transient receptor potential vanilloid

